# OnTheFly^2.0^: a text-mining web application for automated biomedical entity recognition, document annotation, network and functional enrichment analysis

**DOI:** 10.1101/2021.05.14.444150

**Authors:** Fotis A. Baltoumas, Sofia Zafeiropoulou, Evangelos Karatzas, Savvas Paragkamian, Foteini Thanati, Ioannis Iliopoulos, Aristides G. Eliopoulos, Reinhard Schneider, Lars Juhl Jensen, Evangelos Pafilis, Georgios A. Pavlopoulos

## Abstract

Extracting and processing information from documents is of great importance as lots of experimental results and findings are stored in local files. Therefore, extracting and analysing biomedical terms from such files in an automated way is absolutely necessary. In this article, we present OnTheFly^2.0^, a web application for extracting biomedical entities from individual files such as plain texts, Office documents, PDF files or images. OnTheFly^2.0^ can generate informative summaries in popup windows containing knowledge related to the identified terms along with links to various databases. It uses the EXTRACT tagging service to perform Named Entity Recognition (NER) for genes/proteins, chemical compounds, organisms, tissues, environments, diseases, phenotypes and Gene Ontology terms. Multiple files can be analysed, whereas identified terms such as proteins or genes can be explored through functional enrichment analysis or be associated with diseases and PubMed entries. Finally, protein-protein and protein-chemical networks can be generated with the use of STRING and STITCH services. To demonstrate its capacity for knowledge discovery, we interrogated published meta-analyses of clinical biomarkers of severe COVID-19 and uncovered inflammatory and senescence pathways that impact disease pathogenesis. OnTheFly^2.0^ currently supports 197 species and is available at http://onthefly.pavlopouloslab.info.

## INTRODUCTION

The extraction and processing of information from literature is of paramount importance in biomedical sciences. More often than not, researchers need to cope with the task of manually sifting through large amounts of text in various forms (e.g., scientific articles in various file formats, data in spreadsheets and images) in order to obtain pertinent biological information about genes, proteins, chemical compounds, organisms and biological processes and functions. The manual approach is summarized as follows; *i)* read through the article texts, *ii)* detect biomedical entities of interest, and *iii)* query one or more databases for the relevant information. As the volume of literature and experiment-derived datasets continues to increase, this iterative procedure can become slow and heavy. For this reason, text-mining methods are often employed to aid researchers in automatically extracting and searching meaningful biological terms from texts.

One particular application of text mining that is widely used in scientific text processing, is Named Entity Recognition (NER), i.e., identifying words or phrases of interest (the so-called “named entities”) mentioned in plain text, and normalizing them to appropriate database/ontology identifiers (1). In biological and biomedical sciences, these entities include gene and protein names, organisms (scientific or common names), chemical compounds, and ontology terms such as biological processes, cellular components, molecular functions, diseases, phenotypes, and environmental descriptors. Numerous computational tools and web services for NER have been proposed (reviewed in (2–4)). Characteristic examples include tools like EXTRACT (5), PubTator (6), HunFlair (7), BioTextQuest^+^ (8), Saber (9), OGER++ (10) and others. Altogether, these tools detect genes/proteins, genetic variants, diseases, chemical compounds, organisms, diseases and cell lines mentioned in documents. However, while being able to successfully identify entities is critical, NER is only one of the components for meaningful parsing and analysis of the scientific literature. The result of running NER on a set of documents will be a long list of genes and other entities, which the user then needs to navigate and make sense of.

Network visualization is one popular way to get an overview of a large number of entities. Molecular networks can be obtained from a wide variety of different sources, including manually curated pathway databases, e.g., KEGG (11) and Reactome (12), and databases of interaction experiments, e.g., IntAct (13) and BioGRID (14). Resources like STRING (15) and STITCH (16) combine these with additional associations that are predicted or extracted from the biomedical literature through automatic text mining (17). All of these resources can be queried manually by the user, or automatically through Application Programming Interfaces (APIs), packages or plugins. Their results can be used to generate, visualize and analyze interaction networks with network viewers such as Cytoscape (18), Gephi (19), NORMA (20) and others (reviewed (21, 22)).

Functional enrichment analysis is another commonly used approach, which summarizes a long list of genes by comparing it to a collection of gene sets, each representing a Gene Ontology term, molecular pathway, protein domain, disease, etc. Statistically enriched gene sets are then identified by comparing their frequency against a reference background list of genes. Widely used tools for enrichment analysis include DAVID (23), PANTHER (24), WebGestalt (25), aGOtool (26) and g:Profiler (27, 28), each adopting different statistical tests and supporting different enrichment options (reviewed in (29, 30)).

In this article, we present OnTheFly^2.0^, a full, user-friendly pipeline which goes far beyond just applying NER and allows users to start from a collection of documents and, via a set of entities, perform network and enrichment analyses. Through a user-friendly, interactive web interface, OnTheFly^2.0^ supports a plethora of different file formats for text mining and biomedical entity extraction, including text documents (in both editable and read-only formats), spreadsheets and image files. Through NER, OnTheFly^2.0^ can recognize and retrieve a large variety of both biological/biomedical terms. Extracted protein and chemical entities can be combined to create datasets for various types of analyses, including functional enrichment, related literature finding, associations with diseases and protein domain reporting from protein family databases. In addition, OnTheFly^2.0^ allows the generation and visualization of protein-protein and protein-chemical interaction networks. The capabilities of OnTheFly^2.0^ are shown using a case study in which several inflammatory and senescence pathways that impact COVID-19 pathogenesis have been unraveled after analyzing six clinical articles with mentions to clinical biomarkers of severe COVID-19.

## MATERIALS & METHODS

The OnTheFly^2.0^ pipeline consists of four steps (Figure 1): *i)* uploading of input files and conversion from their original format to HTML, *ii)* identification of bioentities with EXTRACT, *iii)* functional annotation on a set of selected identifiers and *iv)* network analysis. A detailed description of these steps is provided in the following subsections.

**Figure 1.**
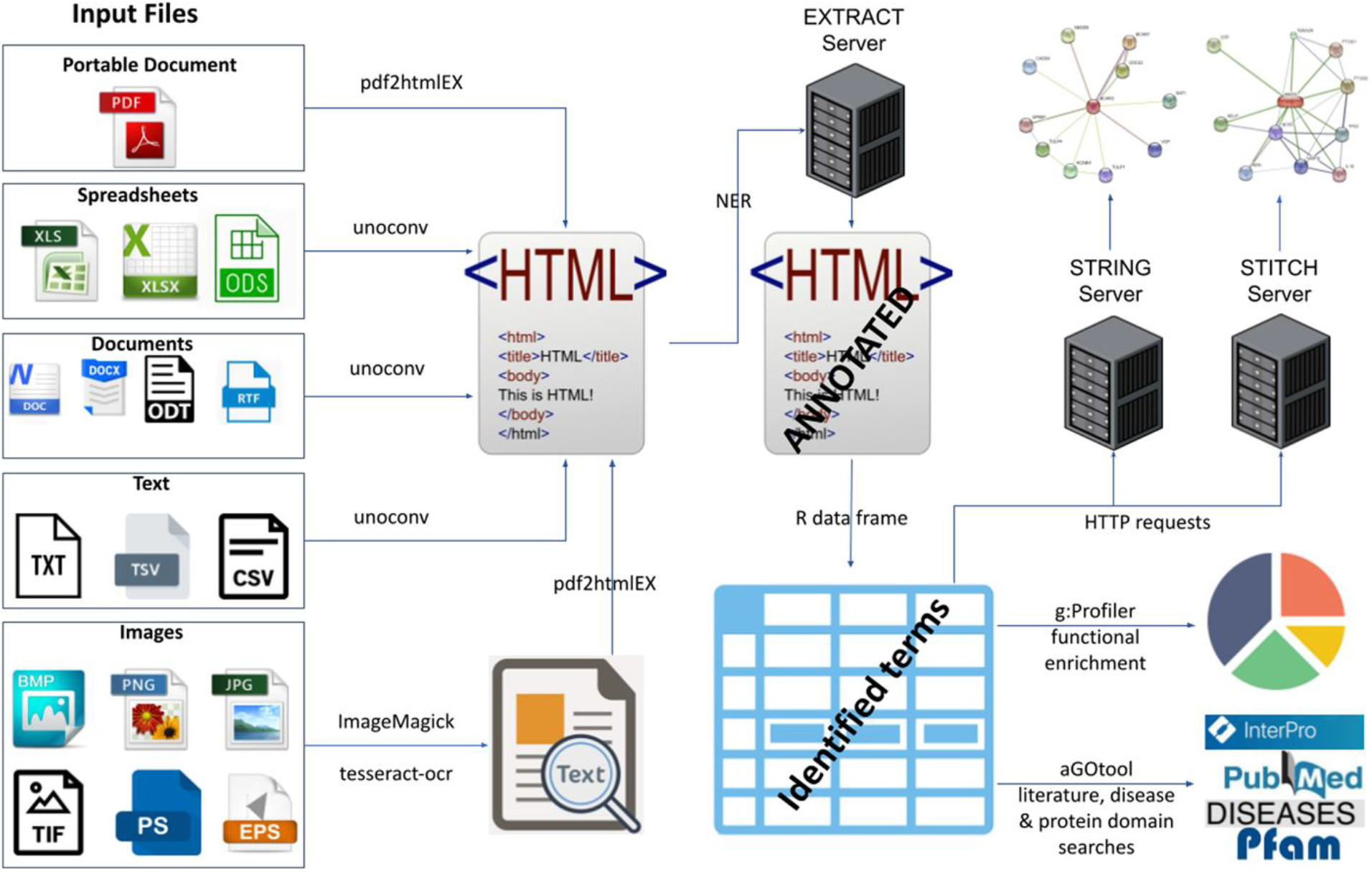
Flowchart of the OnTheFly^2.0^ backend pipeline for file conversion, named entity recognition and data analysis.

### File conversion pipeline

In its current version, OnTheFly^2.0^ supports annotation for PDF files, Office-formatted documents, flat text files and images. In the online version, each file must have a maximum size of 10MBs. Users can upload multiple files simultaneously and process them separately or in combination.

OnTheFly^2.0^ uses various tools and pipelines in its backend to convert uploaded files to HTML format prior to annotation, maintaining overall layout, text formatting, formulas and images to the extent possible. PDF files are converted with the use of *pdf2htmlEX*, an open-source package (31), whereas the *LibreOffice* universal converter (*unoconv*) is used to convert Office files, including formatted/enriched text files (MS Office .doc/.docx, OpenOffice .odt, Rich Text Format .rtf) and spreadsheets (MS Excel .xls/xlsx, OpenOffice Spreadsheet .odp), tab- and comma-delimited table files (.tsv and .csv, respectively) as well as flat text (.txt) files. Notably, in the case of spreadsheet files, the converter is capable of handling each of the sheets.

In addition to the above, OnTheFly^2.0^ can utilize Optical Character Recognition (OCR) scan on images with no text encoding. Both common (.bmp, .jpg, .png, .tiff) and PostScript-compliant image file formats (.ps and .eps) are supported. Image preprocessing and file format conversion is performed with the open-source package *ImageMagick*. OCR scanning of images is performed using the *tesseract-ocr* package (32) to produce PDF files with parseable text, which are then processed as described above. Successful OCR scanning heavily depends on the quality of the imported image, including its resolution. As a result, images containing text elements in rotated orientation or embedded in complex graphical shapes, or images with low resolution may result in poor OCR results.

After files have been uploaded and converted, the resulting HTML version can be inspected in a viewer that has each document in a separate tab. This allows the user to identify conversion or OCR problems before continuing the analysis.

### Document annotation using Named Entity Recognition (NER)

Once the files have been uploaded, users can annotate them with the help of EXTRACT tagging service (5). EXTRACT performs dictionary-based NER using the highly efficient *tagger* software (33) to detect words and phrases, which correspond to biomedical entities. In detail, EXTRACT is capable of identifying environment descriptive terms from Environment Ontology (e.g., desert, forest) (34), organism mentions from NCBI Taxonomy (35), tissue terms from BRENDA Tissue Ontology (36), disease mentions from Disease Ontology (37), phenotypes from Mammalian Phenotype Ontology (38), biological processes, cellular components, molecular functions from Gene Ontology (39, 40), small chemical molecules from PubChem (41), non-coding RNAs from RAIN (42), and protein-coding genes from STRING (15). In the implementation of OnTheFly^2.0^, NER can be performed for a list of 197 organisms.

Once the annotation parameters (entity types and organisms) have been set and a NER process has been completed, OnTheFly^2.0^ will return the annotated document with all of the recognized terms linked and highlighted using different colors (Figure 2). On mouse-click action on a term, OnTheFly^2.0^ will generate a popup window with details about the biomedical entity and links to external databases. In case of term disambiguation (e.g., when a term comes from several organisms or corresponds to more than one entity type), OnTheFly^2.0^ will report all of the possible options. For a more comprehensive summary, all of the identified terms along with their database identifiers and links are collected in an interactive table and can be exported as a CSV file. The table results can be narrowed down after filtering for entity type (e.g., genes/proteins, diseases) at any stage. The annotation process is presented in Figure 2.

**Figure 2.**
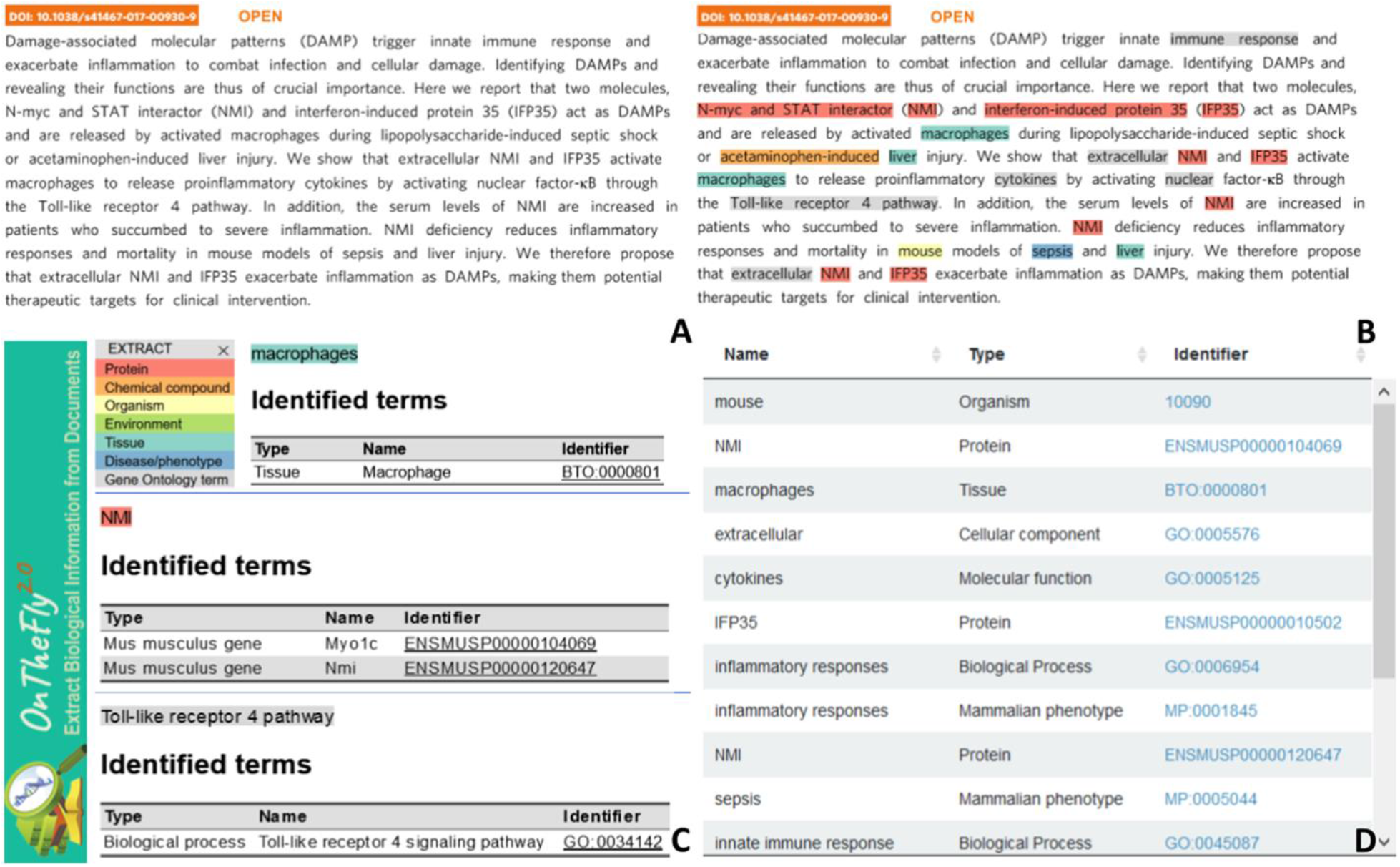
PDF annotation using NER for article by Xiahou et al, 2017 (43). A) The PDF abstract in its simple form. B) The annotated abstract using *M. musculus* as an organism. C) Popup windows with information about the identified term. The term is colored according to its type and original links to external databases are provided. D) A summary table with some of the identified terms.

### Functional enrichment analysis

OnTheFly^2.0^ uses two tools, *g:Profiler* (27, 28) and *aGOtool* (26), to provide rich functional enrichment analysis for a selected set of genes/proteins collected by one or multiple files. The user can customize parameters for the enrichment analysis and choose from a list of 197 organisms. OnTheFly^2.0^ uses g:Profiler to identify enriched functional terms from Gene Ontology (39, 40), pathways from KEGG (11), Reactome (12) and WikiPathways (44), protein complexes from CORUM (45), expression data from Human Protein Atlas (46), regulatory motifs from TRANSFAC (47) and miRTarBase (48), and phenotypes from the Human Phenotype Ontology (49). The analysis results from g:Profiler are complemented by further enrichment analyses from aGOtool to also identify enriched terms from the UniProt keyword classification system, protein families and domains from Pfam (50) and InterPro (51), as well as human diseases from the DISEASES database (52). g:Profiler and aGOtool test for statistically significant enrichment by using Fisher’s exact test to compare the user-defined input dataset (foreground) to a background set from organism-specific genes annotated in the Ensembl database (53) and UniProt Reference Proteomes (54), respectively. The resulting *p*-values are corrected for multiple testing using either g:SCS (only in case of g:Profiler), Bonferroni correction or Benjamini-Hochberg false discovery rate (FDR), all of which can be used as thresholds for the results. Enrichment analysis is performed using ENSEMBL IDs as input, while results can be reported as Entrez, UniProt, EMBL, ENSEMBL and RefSeq gene/protein names/identifiers, based on the user’s choice.

Functional enrichment results are reported in interactive searchable tables displaying details about each functional term. One can expand each row of the table to see which of the identified genes/proteins were found to be associated with the functional term. For example, in the case of a KEGG pathway, one can see how many proteins or genes were found to be related to it and get redirected to the KEGG repository to see the actual schema of the pathway in a static form with all of the detected genes/proteins highlighted. In case of g:Profiler, an interactive Manhattan plot is offered for a clearer overview. In this plot, functional terms are grouped along the x-axis and colored by their data source, whereas the y-axis shows the significance (*p*-value) of each term. Hovering over a data point reveals a tooltip with key information about the functional term. Finally, the most significant functional terms are shown as a bar chart, which the user can customize to show the desired number of terms. All of the aforementioned reports can be exported and saved in various file formats (CSV, XLS, PDF). An overview is shown in Figure 3.

**Figure 3.**
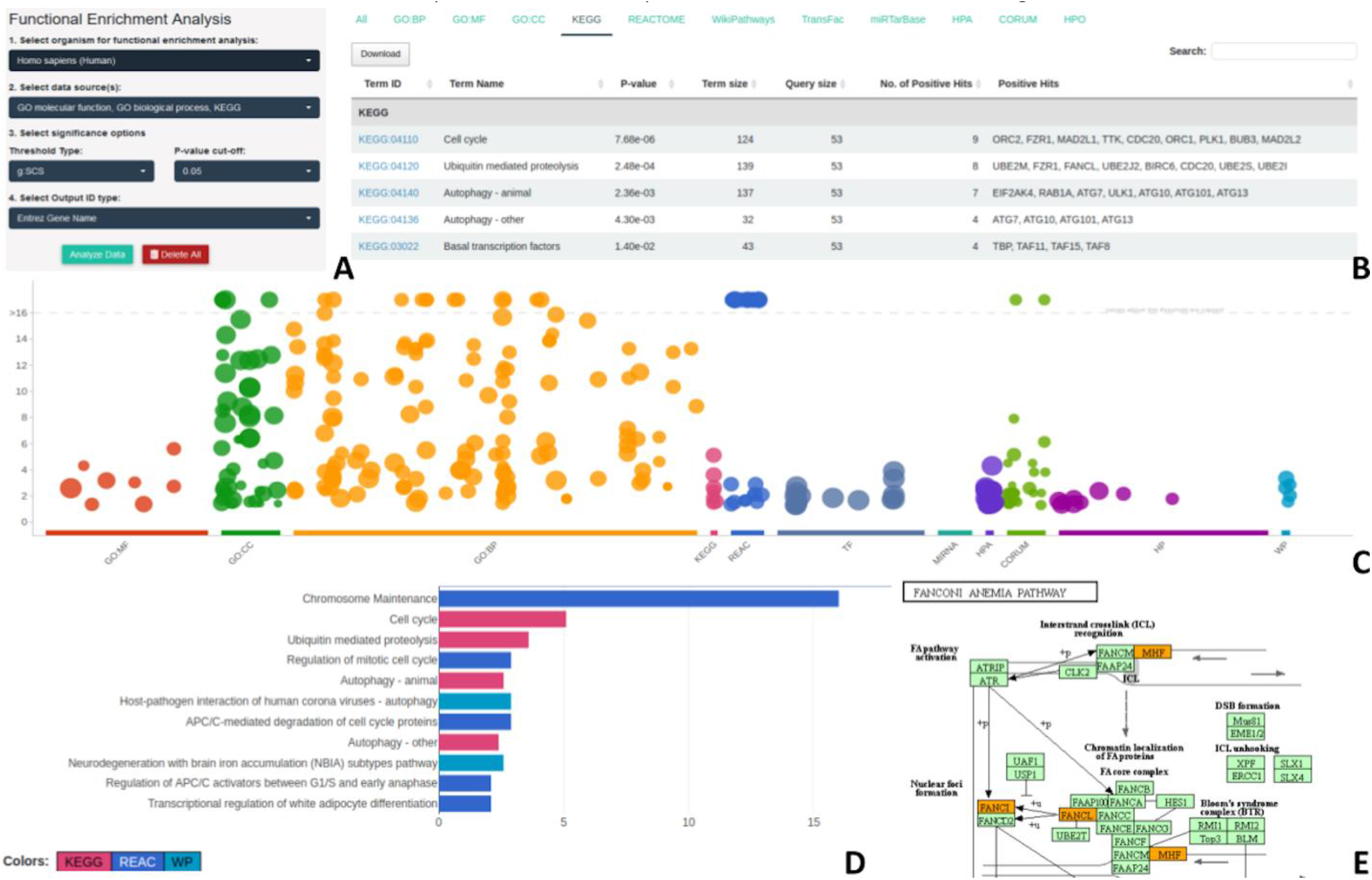
OnTheFly^2.0^’s functional enrichment. A) Functional enrichment input parameters. B) Summary table with the functional terms and the corresponding identified entities. Results from KEGG are shown. C) A functional enrichment overview with the use of a Manhattan plot. D) Bar plot for the distribution of enriched genes into metabolic pathways obtained from three pathway databases (KEGG, Reactome, WikiPathways), with the results of each database colored differently. The bar length is proportional to the extent of enrichment for each term, as represented by -log_10_(P-value). E) Portion of a generated KEGG pathway with the genes found in documents highlighted in orange.

### Publication enrichment analysis

OnTheFly^2.0^ uses the aGOtool to allow users to find scientific articles that mention surprisingly many of the genes/proteins identified in the uploaded input files. While conceptually similar to the functional enrichment analyses just described, publication enrichment analysis serves a very different purpose, namely to help the user identify scientific publications of relevance to the gene/protein list. The publication enrichment analysis in aGOtool is based on a text corpus of all PubMed abstracts and full-text articles from the PubMed Central Open Access subset. These have been run through the same NER *tagger* used in EXTRACT and the results are updated with new documents on a weekly basis. Consequently, all documents have been automatically annotated with the genes mentioned within them, thus turning every document into a gene set. These millions of gene sets are then used by aGOtool in the same manner as all other gene sets.

We make use of this functionality to provide publication enrichment functionality in OnTheFly^2.0^ for the list of 197 organisms. The user can select up to 1,000 of the genes/proteins identified in the uploaded files for analysis, which will then be submitted to aGOtool to test each document from the precomputed corpus for statistically significant enrichment, again using Fisher’s exact test. The resulting *p*-values as well as Bonferroni-corrected *p*-values and Benjamini-Hochberg FDR values can be used for filtering the results. Results are reported in interactive searchable tables displaying details about each literature term (scientific publication or disease). Links are provided for publications and diseases to PubMed. In addition, users are able to rank the most significant publications using barchart plots and manually adjust the number of the reported results with the use of a slide bar. All of the aforementioned reports can be exported and saved in various file formats (CSV, XLS, PDF).

### Network analysis

In addition to the aforementioned enrichment options, OnTheFly^2.0^ offers the capability to construct and visualize biomolecular interaction networks for a set of 197 organisms. This task is performed using the APIs of the STRING (15) and STITCH (16) databases for protein-protein and protein-chemical interactions, respectively. The users may submit their dataset obtained from the uploaded documents to retrieve interactions and visualize the results as networks with the interacting entities presented as nodes and their interactions as edges. For computational efficiency reasons, in its current version, OnTheFly^2.0^ allows a maximum of 500 proteins per request for STRING and 100 proteins or small molecules per request for STITCH.

STRING and STITCH classify interactions between two entities (proteins or small molecules) as either *physical* (i.e., part of the same biomolecular complex), or *functional* (i.e., involved in the same pathway/process). To this end, OnTheFly^2.0^ requires users to select whether to include the *Full* set of interactions (both physical and functional) or the *Physical subnetwork* exclusively. Users can also specify the cutoff on the *Interaction Score*. Finally, users can choose whether each edge should show the type(s) of evidence (e.g., experiments or text mining) supporting it (*Evidence* mode) or if the thickness of the edge should instead show the interaction score (*Confidence* mode).

In addition to the above, in protein-chemical networks network edges can be formatted based on *Molecular Action* or *Binding Affinity*. By choosing *Molecular Action*, the edges in the network will represent the type (activation, inhibition, catalysis etc.) as well as the effect (positive, negative or unspecified) of each protein-chemical interaction. By choosing *Binding Affinity*, the edge thickness will indicate the binding affinity between the proteins and bound chemicals. The resulting network is shown in a separate Network Viewer panel, preserving the characteristic STRING network layout and style. An example of such networks is shown in Suppl. Fig. 1. In addition, options are given to view the generated network in STRING (protein-protein) or STITCH (protein-chemical) for further analysis. Finally, one can export a network as an image or as a tab-delimited file compatible with external network visualization applications.

### Implementation

OnTheFly^2.0^ is a web application implemented in R, using the R/Shiny package as well as HTML, CSS and JavaScript. The Shiny and ShinyJS packages are used as mediators to establish the connection between the R and JavaScript functions. The API of the EXTRACT web service which utilizes the *tagger* text mining utility, is used to perform NER. Functional enrichment analysis is performed using the g:Profiler2 package (R implementation of g:Profiler) and aGOtool. Biological networks are constructed and visualized using the STRING API, as implemented in the STRING and STITCH databases. OnTheFly^2.0^ is available as a web service, and as a standalone package through a GitHub repository. The standalone version is fully functional in native Linux and other Unix-based operating systems. It can also run on Windows, by utilizing a Windows Subsystem for Linux (WSL) or other similar compatibility layers (e.g., Cygwin). The web service is fully functional in all major web browsers (Google Chrome, Mozilla Firefox, Microsoft Edge, Tor, Apple Safari, Opera).

## CASE STUDY

To demonstrate the capacity of OnTheFly^2.0^ for rapid extraction of biological information and knowledge discovery, we analyzed six published meta-analysis reports on clinical biomarkers of severe COVID-19 (55–60) (Suppl. Table 1). Texts in PDF format were annotated by NER, results filtered to manually remove false positives and jointly processed for functional enrichment analysis. Reassuringly, we found “Respiratory failure”, “Pneumonia” and “COVID-19” to be among the most significantly enriched diseases (Suppl. Table 2). The GO enrichment for biological processes (Suppl. Table 3) identified several GO terms related to inflammation, cell activation and response to stress, in line with COVID-19 being associated with exaggerated lung inflammation and systemic immune dysfunction. Similarly, the annotated text terms were found to be enriched for molecular functions that are associated with cytokine activity and cytokine receptor signaling (Suppl. Table 4). These results were supported by the UniProt keyword analysis, which revealed “Cytokine”, “Inflammatory response”, “Host-virus interaction”, and “Host cell receptor for virus entry” to all be enriched (Suppl. Table 5). Analysis of putative protein-protein interactions (physical and functional associations) through the STRING option of OnTheFly^2.0^ uncovered a cluster of interacting cytokines and other immune components that is pertinent to the “cytokine storm” of severe COVID-19 (Figure 4). Cytokines are also a recurring theme in the publication enrichment results, which as one would hope further included several COVID-19 studies (Suppl. Table 6).

**Figure 4.**
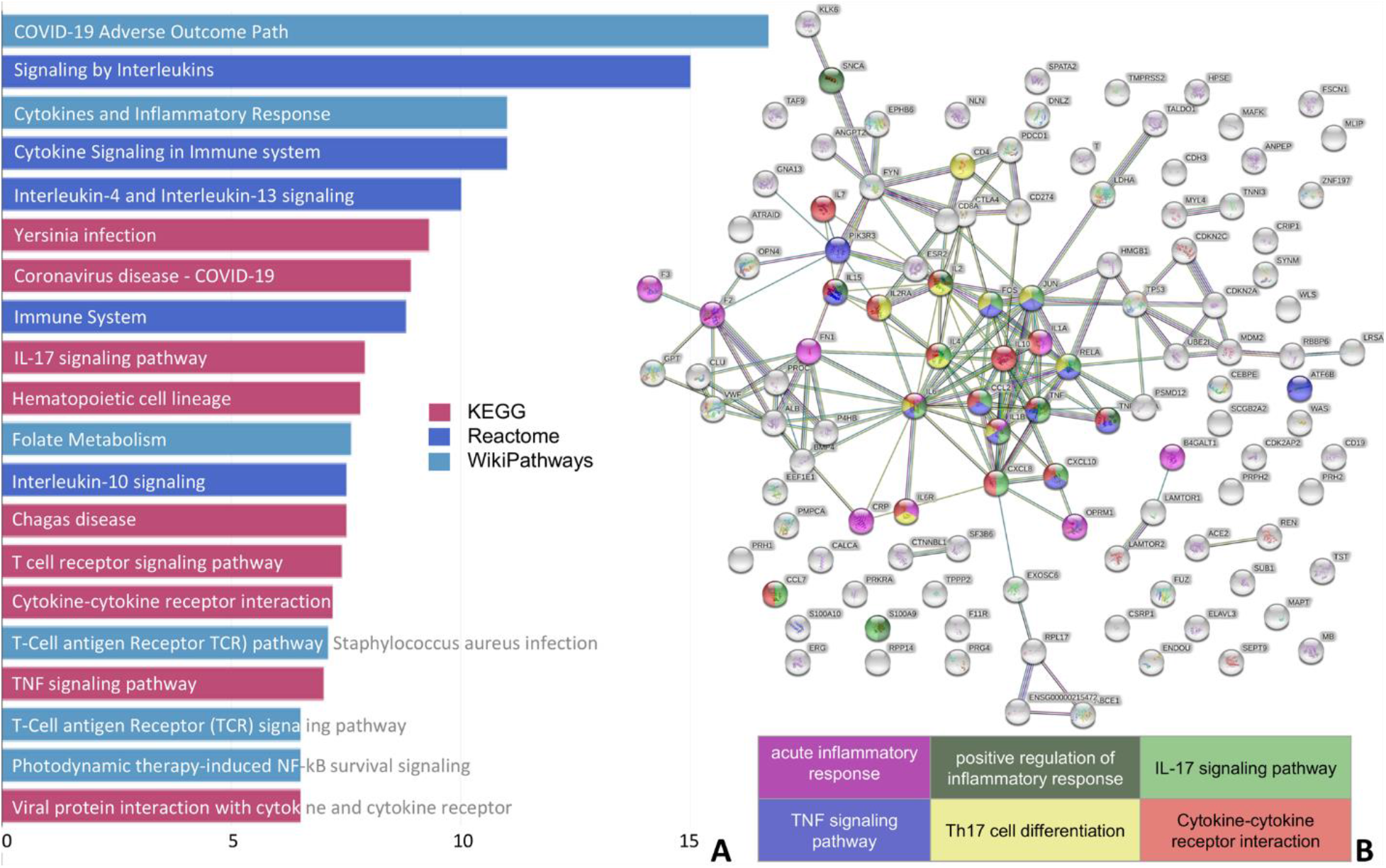
Analysis of clinical biomarkers of severe COVID-19 using OnTheFly^2.0^. A) List of enriched pathways from the KEGG, Reactome and WikiPathways databases. B) Analysis of putative protein-protein interactions through the STRING option of OnTheFly^2.0^. A cluster of interacting components of inflammatory/immune pathways, each represented by a different color, is shown.

Cellular/extracellular components predicted to be associated with biomarkers of severe COVID-19 included extracellular space (GO:0005615, GO:0005576), plasma membrane (GO:0009897, GO:0009986, GO:0098552) and, interestingly, membrane microdomains (also called “membrane rafts”; GO:0098857, GO:0045121) (Suppl. Table 7). The latter emerge as important cellular components implicated in *i)* the initial binding of SARS-CoV-2 to ACE2 receptor, *ii)* virus internalization and *iii)* cell-to-cell transmission (reviewed in (61)). Pertinent to knowledge discovery, this biological information was extracted in the absence of specific reference to membrane microdomains in any of the six meta-analysis reports that were interrogated.

Several relevant KEGG pathways were also extracted (Suppl. Table 8), including “coronavirus disease - COVID-19” (KEGG: 05171; Suppl. Fig. 2), “viral protein interaction with cytokine and cytokine receptor” (KEGG: 04061) and “cytokine-cytokine receptor interaction” (KEGG: 04060). Interestingly, “Yersinia infection” (KEGG: 05135) was also identified as a relevant KEGG pathway with high probability (*p*-value<10^−8^). *Yersinia pestis* is the causative pathogen for pneumonic plague, one of the world’s deadliest infectious diseases. *Yersinia pestis* infects pneumocytes and alveolar macrophages, triggering inflammasome-mediated IL-1β/IL-18 cytokine release (62) that is followed by neutrophil influx, exaggerated inflammation and lung tissue damage (63). These immune and lung tissue reactions to *Yersinia pestis* are reminiscent of those to severe SARS-CoV-2 infection (61) and warrant further insights into the immunological mechanisms of response to these unrelated pathogens. Of additional interest is the predicted involvement of the “IL-17 signaling pathway” (KEGG: 04657) in severe COVID-19 (Suppl. Table 8) which is supported by a recent study reporting T cell skewing towards Th17, a specialized CD4^+^ effector T cell lineage characterized by secretion of IL-17 and IL-17F cytokines in patients with COVID-19 pneumonia (64).

We also explored the REACTOME option of OnTheFly^2.0^ to map and analyze biological pathways that are over-represented in the validation example. As shown in Suppl. Table 9, several cytokine pathways were predicted to be significantly associated with biomarkers of severe COVID-19. We note that predicted REACTOME pathways included “cellular senescence” despite the absence of specific references to this biological term in any of the six annotated meta-analysis reports under study. In line with this prediction, COVID-19 pneumonia has recently been associated with immunosenescence (64) and accelerated aging of pneumocytes (65). Overall, the aforementioned analyses underscore the practical utility of OnTheFly^2.0^ to rapidly extract biological information from texts and hence assisting knowledge discovery (Figure 4).

## DISCUSSION

OnTheFly^2.0^ has been redeveloped to use current technologies and overcome many of the problems of its predecessor (66). The GUI has been completely rewritten to no longer rely on a Java applet and instead using R, Shiny, CSS, HTML and JavaScript technologies. The backend document format conversion has also been considerably improved, replacing commercial Windows-based converters with open-source, Unix-based ones, which furthermore do a much better job preserving the original document layout. Moreover, compared to its predecessor, OnTheFly^2.0^ comes with a broader spectrum of term types it can identify and supports OCR technology for processing images. Uploaded files are only stored temporarily in the OnTheFly^2.0^ server just for parsing and no file backups, copies or personal data are kept. A more detailed comparison between OnTheFly^1.0^ and OnTheFly^2.0^ is presented in Table 1.

**Table 1.** OnTheFly^1.0^ vs OnTheFly^2.0^

| Functionality | OnTheFly<sup>1.0</sup> | OnTheFly<sup>2.0</sup> |
| --- | --- | --- |
| Named Entity Recognition | genes, proteins, chemical compounds | genes/proteins, chemical compounds, organism names, environments, tissues, diseases, phenotypes, gene ontology terms |
| Supported files in textual formats | PDF (.pdf),<br>Office texts (.doc),<br>Flat text (.txt, .tsv, .csv) | PDF (.pdf)<br>Microsoft Word (.doc and .docx)<br>OpenOffice Writer(.odt)<br>Microsoft Excel (.xls, .xlsm and .xlsx)<br>OpenOffice Calc (.ods)<br>Flat text (.txt, .tsv, .csv) |
| Image files with the use of OCR | N/A | Images (.bmp, .png, .jpg, .tif),<br>PostScript (.ps, .eps)<br>*tesseract-ocr package* |
| Interaction networks | STRING | STRING, STITCH |
| Infrastructure | Windows-based server | Unix-based (Linux) server and<br>standalone package, compatible with<br>Windows through WSL (Windows<br>Subsystem for Linux) |
| File Converters | Commercial converters:<br>ultrashareware, verypdf<br>(PDF layout loss) | Freely available, open-source<br>converters:<br>pdf2htmlEX, LibreOffice,<br>ImageMagick<br>(PDF, Text, spreadsheet & image<br>layout preservation with minimal<br>losses) |
| Tagging service | REFLECT | EXTRACT |
| GUI Implementation | Java Applet (obsolete) | R/Shiny, JavaScript |
| Functional Enrichment | Link to<br>BioCompendium (outdated and<br>Human/Mouse only) | In-house analysis based on g:Profiler<br>and aGOTool. Enrichment terms<br>include Gene Ontology, biological<br>pathways, regulatory motifs, protein<br>databases, human phenotype<br>ontology etc. |
| Organisms | Human and Mouse | 197 species |
| Combination of files and entity<br>selection | N/A | Offered |
| Parameterization | N/A | Offered (e.g., functional enrichment<br>options) |
| Browser Compatibility | Java Applets are not supported<br>anymore | Safari, Tor, Firefox, Chrome, Edge,<br>Opera |

## CONCLUSIONS

OnTheFly^2.0^ is a powerful tool for identifying terms in locally stored documents varying from texts and PDFs to Office and image files. Users can identify terms such as proteins, genes, chemical compounds, organisms, tissues, environments, diseases, phenotypes and gene ontologies and perform a functional enrichment and network analysis upon selecting a set of biomedical entities. Furthermore, popup windows with informative summaries about a term and its links to external repositories are also generated. OnTheFly^2.0^ can aid researchers in annotating locally stored documents and further exploring and analysing their identified biomedical entities in a fully automated way. We believe that due to its offered capabilities and ease of use, OnTheFly^2.0^ will reach a broad spectrum of users varying from experimentalists to bioinformaticians.

## AVAILABILITY

OnTheFly^2.0^ is available at: http://onthefly.pavlopouloslab.info or http://bib.fleming.gr:3838/OnTheFly/.

The source code and instructions about the necessary dependencies can be found at https://github.com/PavlopoulosLab/OnTheFly.

## REFERENCES

1. Nadeaux, D. and Sekine, S. (2007) A survey of named entity recognition and classification. Lingvisticæ Investigationes, 30, 3–26.

2. Rebholz-Schuhmann, D., Oellrich, A. and Hoehndorf, R. (2012) Text-mining solutions for biomedical research: enabling integrative biology. Nat Rev Genet, 13, 829–839.

3. Przybyła, P., Shardlow, M., Aubin, S., Bossy, R., Eckart de Castilho, R., Piperidis, S., McNaught, J. and Ananiadou, S. (2016) Text mining resources for the life sciences. Database (Oxford), 2016.

4. Perera, N., Dehmer, M. and Emmert-Streib, F. (2020) Named Entity Recognition and Relation Detection for Biomedical Information Extraction. Front Cell Dev Biol, 8, 673.

5. Pafilis, E., Buttigieg, P.L., Ferrell, B., Pereira, E., Schnetzer, J., Arvanitidis, C. and Jensen, L.J. (2016) EXTRACT: interactive extraction of environment metadata and term suggestion for metagenomic sample annotation. Database (Oxford), 2016.

6. Wei, C.-H., Kao, H.-Y. and Lu, Z. (2013) PubTator: a web-based text mining tool for assisting biocuration. Nucleic Acids Res, 41, W518–522.

7. Weber, L., Sänger, M., Münchmeyer, J., Habibi, M., Leser, U. and Akbik, A. (2021) HunFlair: An Easy-to-Use Tool for State-of-the-Art Biomedical Named Entity Recognition. Bioinformatics, 10.1093/bioinformatics/btab042.

8. Papanikolaou, N., Pavlopoulos, G.A., Pafilis, E., Theodosiou, T., Schneider, R., Satagopam, V.P., Ouzounis, C.A., Eliopoulos, A.G., Promponas, V.J. and Iliopoulos, I. (2014) BioTextQuest(+): a knowledge integration platform for literature mining and concept discovery. Bioinformatics, 30, 3249–3256.

9. Giorgi, J.M. and Bader, G.D. (2020) Towards reliable named entity recognition in the biomedical domain. Bioinformatics, 36, 280–286.

10. Furrer, L., Jancso, A., Colic, N. and Rinaldi, F. (2019) OGER++: hybrid multi-type entity recognition. J Cheminform, 11, 7.

11. Kanehisa, M., Furumichi, M., Sato, Y., Ishiguro-Watanabe, M. and Tanabe, M. (2021) KEGG: integrating viruses and cellular organisms. Nucleic Acids Res, 49, D545–D551.

12. Jassal, B., Matthews, L., Viteri, G., Gong, C., Lorente, P., Fabregat, A., Sidiropoulos, K., Cook, J., Gillespie, M., Haw, R., et al. (2020) The reactome pathway knowledgebase. Nucleic Acids Res, 48, D498–D503.

13. Orchard, S., Ammari, M., Aranda, B., Breuza, L., Briganti, L., Broackes-Carter, F., Campbell, N.H., Chavali, G., Chen, C., del-Toro, N., et al. (2014) The MIntAct project--IntAct as a common curation platform for 11 molecular interaction databases. Nucleic Acids Res, 42, D358–363.

14. Oughtred, R., Rust, J., Chang, C., Breitkreutz, B.-J., Stark, C., Willems, A., Boucher, L., Leung, G., Kolas, N., Zhang, F., et al. (2021) The BioGRID database: A comprehensive biomedical resource of curated protein, genetic, and chemical interactions. Protein Sci, 30, 187–200.

15. Szklarczyk, D., Gable, A.L., Nastou, K.C., Lyon, D., Kirsch, R., Pyysalo, S., Doncheva, N.T., Legeay, M., Fang, T., Bork, P., et al. (2021) The STRING database in 2021: customizable protein–protein networks, and functional characterization of user-uploaded gene/measurement sets. Nucleic Acids Research, 49, D605–D612.

16. Szklarczyk, D., Santos, A., von Mering, C., Jensen, L.J., Bork, P. and Kuhn, M. (2016) STITCH 5: augmenting protein-chemical interaction networks with tissue and affinity data. Nucleic Acids Res, 44, D380–384.

17. Koutrouli, M., Hatzis, P. and Pavlopoulos, G.A. (2021) Exploring Networks in the STRING and Reactome Database. In Wolkenhauer, O. (ed), Systems Medicine. Academic Press, Oxford, pp. 507–520.

18. Shannon, P., Markiel, A., Ozier, O., Baliga, N.S., Wang, J.T., Ramage, D., Amin, N., Schwikowski, B. and Ideker, T. (2003) Cytoscape: a software environment for integrated models of biomolecular interaction networks. Genome Res, 13, 2498–2504.

19. Bastian, M., Heymann, S. and Jacomy, M. (2009) Gephi: An Open Source Software for Exploring and Manipulating Networks.

20. Koutrouli, M., Karatzas, E., Papanikolopoulou, K. and Pavlopoulos, G.A. (2020) NORMA-The network makeup artist: a web tool for network annotation visualization. bioRxiv, 10.1101/2020.03.05.978585.

21. Pavlopoulos, G.A., Wegener, A.-L. and Schneider, R. (2008) A survey of visualization tools for biological network analysis. BioData Min, 1, 12.

22. Koutrouli, M., Karatzas, E., Paez-Espino, D. and Pavlopoulos, G.A. (2020) A Guide to Conquer the Biological Network Era Using Graph Theory. Front Bioeng Biotechnol, 8, 34.

23. Jiao, X., Sherman, B.T., Huang, D.W., Stephens, R., Baseler, M.W., Lane, H.C. and Lempicki, R.A. (2012) DAVID-WS: a stateful web service to facilitate gene/protein list analysis. Bioinformatics, 28, 1805–1806.

24. Mi, H., Ebert, D., Muruganujan, A., Mills, C., Albou, L.-P., Mushayamaha, T. and Thomas, P.D. (2021) PANTHER version 16: a revised family classification, tree-based classification tool, enhancer regions and extensive API. Nucleic Acids Res, 49, D394–D403.

25. Liao, Y., Wang, J., Jaehnig, E.J., Shi, Z. and Zhang, B. (2019) WebGestalt 2019: gene set analysis toolkit with revamped UIs and APIs. Nucleic Acids Res, 47, W199–W205.

26. Schölz, C., Lyon, D., Refsgaard, J.C., Jensen, L.J., Choudhary, C. and Weinert, B.T. (2015) Avoiding abundance bias in the functional annotation of post-translationally modified proteins. Nat Methods, 12, 1003–1004.

27. Raudvere, U., Kolberg, L., Kuzmin, I., Arak, T., Adler, P., Peterson, H. and Vilo, J. (2019) g:Profiler: a web server for functional enrichment analysis and conversions of gene lists (2019 update). Nucleic Acids Res, 47, W191–W198.

28. Kolberg, L., Raudvere, U., Kuzmin, I., Vilo, J. and Peterson, H. (2020) gprofiler2 --an R package for gene list functional enrichment analysis and namespace conversion toolset g:Profiler. F1000Res, 9.

29. Maleki, F., Ovens, K., Hogan, D.J. and Kusalik, A.J. (2020) Gene Set Analysis: Challenges, Opportunities, and Future Research. Front Genet, 11, 654.

30. Mathur, R., Rotroff, D., Ma, J., Shojaie, A. and Motsinger-Reif, A. (2018) Gene set analysis methods: a systematic comparison. BioData Min, 11, 8.

31. Wang, L. and Liu, W. Online publishing via pdf2htmlEX. In TUGboat. Tex Users Group, Tokyo, Japan, Vol. 34 No. 3, pp. 313–324.

32. Smith, R. (2007) An Overview of the Tesseract OCR Engine. In Ninth International Conference on Document Analysis and Recognition (ICDAR 2007).Vol. 2, pp. 629–633.

33. Pafilis, E. and Jensen, L.J. (2016) Real-time tagging of biomedical entities. bioRxiv, 10.1101/078469.

34. Buttigieg, P.L., Morrison, N., Smith, B., Mungall, C.J., Lewis, S.E., and ENVO Consortium (2013) The environment ontology: contextualising biological and biomedical entities. J Biomed Semantics, 4, 43.

35. Schoch, C.L., Ciufo, S., Domrachev, M., Hotton, C.L., Kannan, S., Khovanskaya, R., Leipe, D., Mcveigh, R., O’Neill, K., Robbertse, B., et al. (2020) NCBI Taxonomy: a comprehensive update on curation, resources and tools. Database (Oxford), 2020.

36. Gremse, M., Chang, A., Schomburg, I., Grote, A., Scheer, M., Ebeling, C. and Schomburg, D. (2011) The BRENDA Tissue Ontology (BTO): the first all-integrating ontology of all organisms for enzyme sources. Nucleic Acids Res, 39, D507–513.

37. Schriml, L.M., Mitraka, E., Munro, J., Tauber, B., Schor, M., Nickle, L., Felix, V., Jeng, L., Bearer, C., Lichenstein, R., et al. (2019) Human Disease Ontology 2018 update: classification, content and workflow expansion. Nucleic Acids Res, 47, D955–D962.

38. Smith, C.L. and Eppig, J.T. (2009) The mammalian phenotype ontology: enabling robust annotation and comparative analysis. Wiley Interdiscip Rev Syst Biol Med, 1, 390–399.

39. Ashburner, M., Ball, C.A., Blake, J.A., Botstein, D., Butler, H., Cherry, J.M., Davis, A.P., Dolinski, K., Dwight, S.S., Eppig, J.T., et al. (2000) Gene ontology: tool for the unification of biology. The Gene Ontology Consortium. Nat Genet, 25, 25–29.

40. Gene Ontology Consortium (2021) The Gene Ontology resource: enriching a GOld mine. Nucleic Acids Res, 49, D325–D334.

41. Kim, S., Chen, J., Cheng, T., Gindulyte, A., He, J., He, S., Li, Q., Shoemaker, B.A., Thiessen, P.A., Yu, B., et al. (2021) PubChem in 2021: new data content and improved web interfaces. Nucleic Acids Res, 49, D1388–D1395.

42. Junge, A., Refsgaard, J.C., Garde, C., Pan, X., Santos, A., Alkan, F., Anthon, C., von Mering, C., Workman, C.T., Jensen, L.J., et al. (2017) RAIN: RNA-protein Association and Interaction Networks. Database (Oxford), 2017.

43. Xiahou, Z., Wang, X., Shen, J., Zhu, X., Xu, F., Hu, R., Guo, D., Li, H., Tian, Y., Liu, Y., et al. (2017) NMI and IFP35 serve as proinflammatory DAMPs during cellular infection and injury. Nat Commun, 8, 950.

44. Martens, M., Ammar, A., Riutta, A., Waagmeester, A., Slenter, D.N., Hanspers, K. A Miller, R., Digles, D., Lopes, E.N., Ehrhart, F., et al. (2021) WikiPathways: connecting communities. Nucleic Acids Res, 49, D613–D621.

45. Giurgiu, M., Reinhard, J., Brauner, B., Dunger-Kaltenbach, I., Fobo, G., Frishman, G., Montrone, C. and Ruepp, A. (2019) CORUM: the comprehensive resource of mammalian protein complexes-2019. Nucleic Acids Res, 47, D559–D563.

46. Uhlén, M., Fagerberg, L., Hallström, B.M., Lindskog, C., Oksvold, P., Mardinoglu, A., Sivertsson, Å., Kampf, C., Sjöstedt, E., Asplund, A., et al. (2015) Proteomics. Tissue-based map of the human proteome. Science, 347, 1260419.

47. Wingender, E. (2008) The TRANSFAC project as an example of framework technology that supports the analysis of genomic regulation. Brief Bioinform, 9, 326–332.

48. Huang, H.-Y., Lin, Y.-C.-D., Li, J., Huang, K.-Y., Shrestha, S., Hong, H.-C., Tang, Y., Chen, Y.-G., Jin, C.-N., Yu, Y., et al. (2020) miRTarBase 2020: updates to the experimentally validated microRNA-target interaction database. Nucleic Acids Res, 48, D148–D154.

49. Köhler, S., Gargano, M., Matentzoglu, N., Carmody, L.C., Lewis-Smith, D., Vasilevsky, N.A., Danis, D., Balagura, G., Baynam, G., Brower, A.M., et al. (2021) The Human Phenotype Ontology in 2021. Nucleic Acids Res, 49, D1207–D1217.

50. Mistry, J., Chuguransky, S., Williams, L., Qureshi, M., Salazar, G.A., Sonnhammer, E.L.L., Tosatto, S.C.E., Paladin, L., Raj, S., Richardson, L.J., et al. (2021) Pfam: The protein families database in 2021. Nucleic Acids Research, 49, D412–D419.

51. Blum, M., Chang, H.-Y., Chuguransky, S., Grego, T., Kandasaamy, S., Mitchell, A., Nuka, G., Paysan-Lafosse, T., Qureshi, M., Raj, S., et al. (2021) The InterPro protein families and domains database: 20 years on. Nucleic Acids Res, 49, D344–D354.

52. Pletscher-Frankild, S., Pallejà, A., Tsafou, K., Binder, J.X. and Jensen, L.J. (2015) DISEASES: text mining and data integration of disease-gene associations. Methods, 74, 83–89.

53. Howe, K.L., Achuthan, P., Allen, J., Allen, J., Alvarez-Jarreta, J., Amode, M.R., Armean, I.M., Azov, A.G., Bennett, R., Bhai, J., et al. (2021) Ensembl 2021. Nucleic Acids Research, 49, D884–D891.

54. The UniProt Consortium (2021) UniProt: the universal protein knowledgebase in 2021. Nucleic Acids Research, 49, D480–D489.

55. Henry, B.M., de Oliveira, M.H.S., Benoit, S., Plebani, M. and Lippi, G. (2020) Hematologic, biochemical and immune biomarker abnormalities associated with severe illness and mortality in coronavirus disease 2019 (COVID-19): a meta-analysis. Clin Chem Lab Med, 58, 1021–1028.

56. Danwang, C., Endomba, F.T., Nkeck, J.R., Wouna, D.L.A., Robert, A. and Noubiap, J.J. (2020) A meta-analysis of potential biomarkers associated with severity of coronavirus disease 2019 (COVID-19). Biomark Res, 8, 37.

57. Leisman, D.E., Ronner, L., Pinotti, R., Taylor, M.D., Sinha, P., Calfee, C.S., Hirayama, A.V., Mastroiani, F., Turtle, C.J., Harhay, M.O., et al. (2020) Cytokine elevation in severe and critical COVID-19: a rapid systematic review, meta-analysis, and comparison with other inflammatory syndromes. Lancet Respir Med, 8, 1233–1244.

58. Elshazli, R.M., Toraih, E.A., Elgaml, A., El-Mowafy, M., El-Mesery, M., Amin, M.N., Hussein, M.H., Killackey, M.T., Fawzy, M.S. and Kandil, E. (2020) Diagnostic and prognostic value of hematological and immunological markers in COVID-19 infection: A meta-analysis of 6320 patients. PLoS One, 15, e0238160.

59. Figliozzi, S., Masci, P.G., Ahmadi, N., Tondi, L., Koutli, E., Aimo, A., Stamatelopoulos, K., Dimopoulos, M.-A., Caforio, A.L.P. and Georgiopoulos, G. (2020) Predictors of adverse prognosis in COVID-19: A systematic review and meta-analysis. Eur J Clin Invest, 50, e13362.

60. Tian, W., Jiang, W., Yao, J., Nicholson, C.J., Li, R.H., Sigurslid, H.H., Wooster, L., Rotter, J.I., Guo, X. and Malhotra, R. (2020) Predictors of mortality in hospitalized COVID-19 patients: A systematic review and meta-analysis. J Med Virol, 92, 1875–1883.

61. Gkouskou, K., Vasilogiannakopoulou, T., Andreakos, E., Davanos, N., Gazouli, M., Sanoudou, D. and Eliopoulos, A.G. (2021) COVID-19 enters the expanding network of apolipoprotein E4-related pathologies. Redox Biol, 41, 101938.

62. Sivaraman, V., Pechous, R.D., Stasulli, N.M., Eichelberger, K.R., Miao, E.A. and Goldman, W.E. (2015) Yersinia pestis activates both IL-1β and IL-1 receptor antagonist to modulate lung inflammation during pneumonic plague. PLoS Pathog, 11, e1004688.

63. Pechous, R.D., Sivaraman, V., Price, P.A., Stasulli, N.M. and Goldman, W.E. (2013) Early host cell targets of Yersinia pestis during primary pneumonic plague. PLoS Pathog, 9, e1003679.

64. De Biasi, S., Meschiari, M., Gibellini, L., Bellinazzi, C., Borella, R., Fidanza, L., Gozzi, L., Iannone, A., Lo Tartaro, D., Mattioli, M., et al. (2020) Marked T cell activation, senescence, exhaustion and skewing towards TH17 in patients with COVID-19 pneumonia. Nat Commun, 11, 3434.

65. Evangelou, K., Veroutis, D., Foukas, P.G., Paschalaki, K., Kittas, C., Tzioufas, A.G., Leval, L. de, Vassilakos, D., Barnes, P.J. and Gorgoulis, V.G. (2021) Alveolar type II cells harbouring SARS-CoV-2 show senescence with a proinflammatory phenotype. bioRxiv, 10.1101/2021.01.02.424917.

66. Pavlopoulos, G.A., Pafilis, E., Kuhn, M., Hooper, S.D. and Schneider, R. (2009) OnTheFly: a tool for automated document-based text annotation, data linking and network generation. Bioinformatics, 25, 977–978.

