## Supplementary figures and images for "OnTheFly^2.0^: a text-mining web application for automated biomedical entity recognition, document annotation, network and functional enrichment analysis"

### Suppl_Figure_1.tif

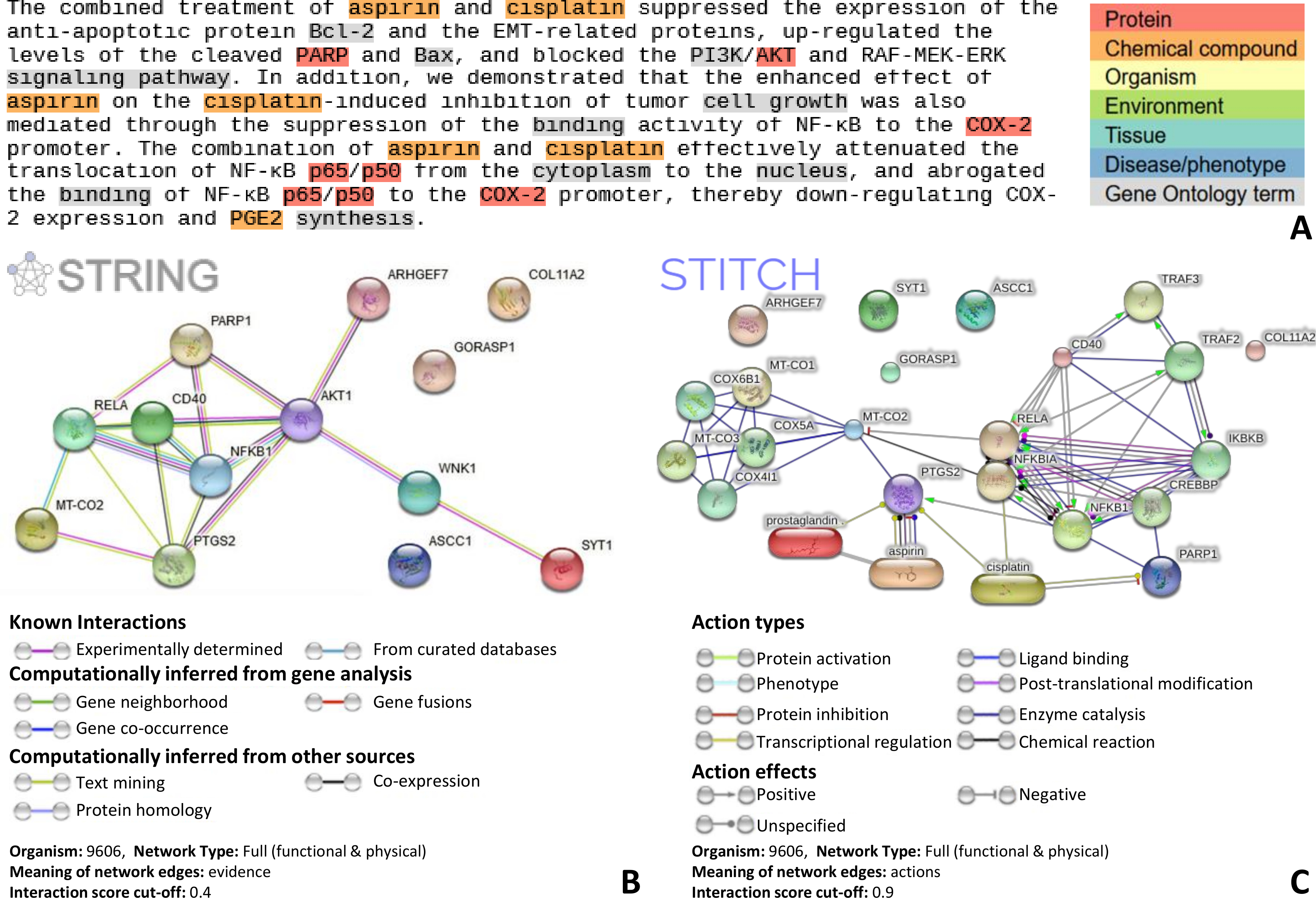

### Suppl_Figure_2.tif

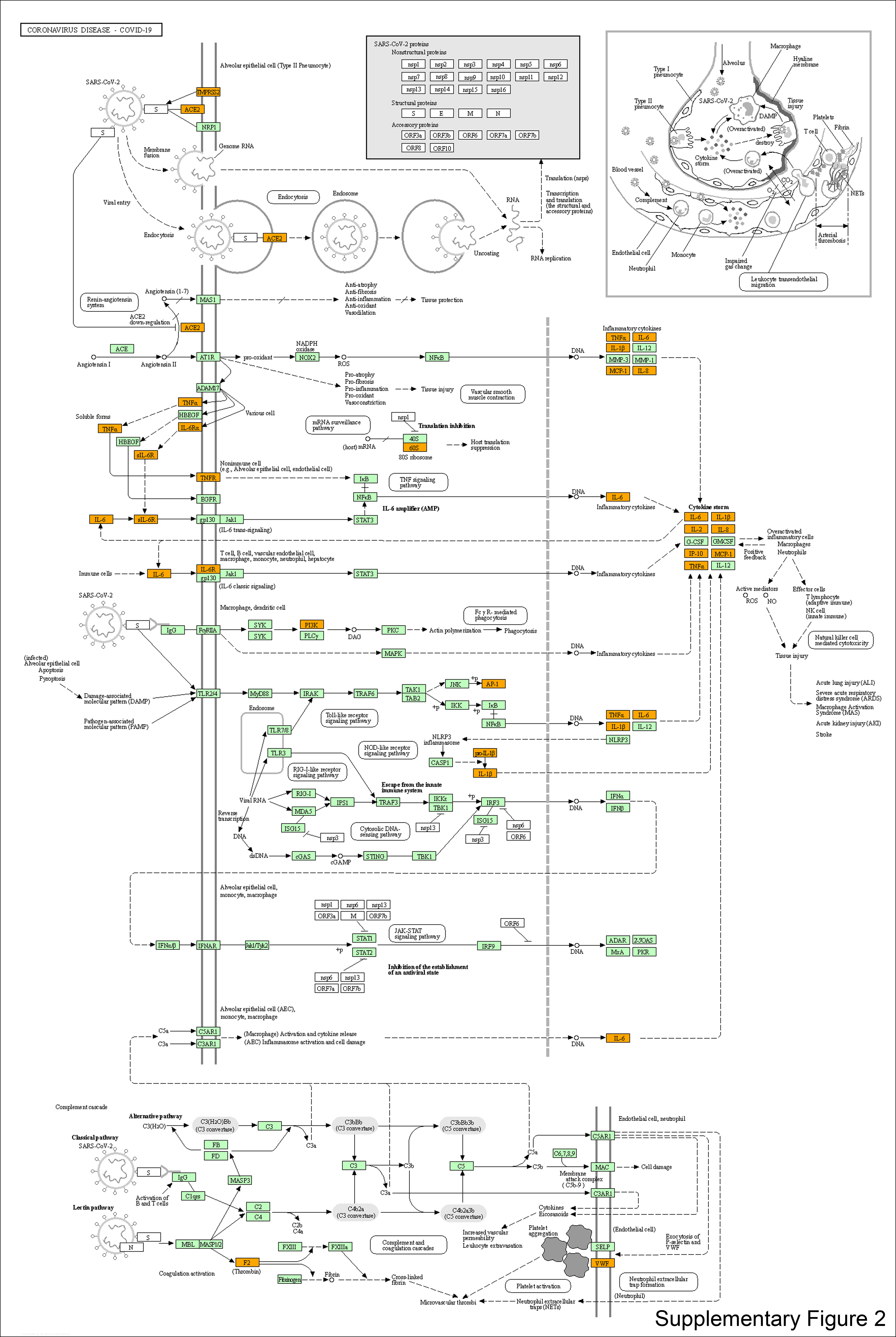
